# Distinct pools of CENP-A and CENP-C support unique phases of centromere assembly in spermatogenesis

**DOI:** 10.64898/2026.01.07.698108

**Authors:** Rachel S Keegan, Elaine M Dunleavy

## Abstract

The centromere is the chromosomal site of kinetochore assembly, defined by the histone H3 variant CENP-A. Each cell cycle, the assembly and maintenance of CENP-A is functionally critical for chromosome segregation. In *Drosophila* male meiosis, CID (fly CENP-A) is assembled in two phases; prophase of meiosis I and post-meiosis II. Here we investigate the dynamics of the assembly components CAL1 and CENP-C in prophase I and determine the requirements for the second assembly phase. In early prophase I, CENP-C functions with CAL1 to maintain the centromere. In late prophase I, CAL1 is undetectable at centromeres and CENP-C no longer functions in centromere maintenance. Instead CENP-C is critical for meiotic kinetochore recruitment and function. This CENP-C pool also functions in CID assembly post-meiosis II, which is independent of CAL1. In addition to different functional pools of CENP-C, distinct pools of CID protein persist in the male germline, and the synthesis of each pool is uncoupled from its cell cycle deposition timing.

## Introduction

Accurate segregation of replicated chromosomes is imperative for preserving genomic integrity and preventing aneuploidy. This is especially critical in the germline, where chromosome segregation during gametogenesis ensures the transmission of genetic information to future generations. Centromeres are unique chromosomal loci that are required for faithful chromosome segregation. They serve as platforms for kinetochore assembly, to which spindle microtubules attach during cell division (Kixmoeller et al., 2020; Fukagawa and Earnshaw, 2014). This locus is epigenetically specified by the incorporation of the histone H3 variant known as Centromere Protein A or CENP-A (McKinley and Cheeseman, 2016; Allshire and Karpen, 2008). During DNA replication in S-phase, parental histones are diluted to half on each DNA strand and are replenished with newly synthesised histones at the replication fork. At centromeres, CENP-A is also diluted during DNA replication, but newly synthesised CENP-A does not load with the canonical histones during S-phase (Sundararajan and Straight, 2022). Instead, cells complete mitosis with only half the maximal occupancy of CENP-A nucleosomes, and new deposition occurs between late telophase and early G1-phase (Jansen et al., 2007; Schuh et al., 2007).

*Drosophila* melanogaster, or the fruit fly, presents a simplified centromere composition, containing CENP-A (referred to as CID in flies), Chromosome Alignment Defect 1 (CAL1) and the inner kinetochore protein CENP-C (Kyriacou and Heun, 2023). In *Drosophila*, CENP-C is the only member of the multiprotein constitutive centromere-associated network (CCAN) complex existing in mammals (Foltz et al., 2006; Okada et al., 2006), and it bridges the centromere and the outer kinetochore (Przewloka et al., 2011). CAL1 acts as the CID-specific chaperone and assembly factor (Chen et al., 2014), fulfilling the role of both Holliday Junction Recognising Protein (HJURP) and the Mis18 complex in mammals (Rowley and Jansen, 2025). In mitosis, CID, CAL1, and CENP-C are interdependent for their localisation to the centromere (Erhardt et al., 2008). CID and CAL1 form a pre-nucleosomal complex that is recruited to the centromere by CENP-C. CAL1 recruits CENP-C, which in turn brings more pre-nucleosomal complexes of CID-CAL1, sustaining this epigenetic loop (Schittenhelm et al., 2010; Chen et al., 2014; Roure et al., 2019; Medina-Pritchard et al., 2020).

The mechanism of CENP-A deposition in meiosis, the specialised division in which homologous chromosomes separate to generate haploid gametes, is less well studied than in mitosis. The timing of cell cycle deposition is markedly different based on initial observations from diverse models. In female meiosis, CENP-A is assembled during, or prior to, prophase I in worms (*Caenorhabditis elegans*), starfish (*Patricia miniate*), mice (*Mus musculus*) and *Drosophila* (Monen et al., 2005; Swartz et al., 2019; Smoak et al., 2016; Tower et al., 2025; Fellmeth et al., 2023). In the *Drosophila* male germline, new CID assembly occurs at two distinct windows: initially, during meiotic prophase I and later, immediately after meiosis II (Dunleavy et al., 2012; Raychaudhuri et al., 2012). A recent study indicates that CENP-A assembly in prophase I is conserved in mammalian spermatogenesis (Štiavnická et al., 2025). In *Drosophila* males, both CAL1 and CENP-C are required for the first meiotic loading event, yet both proteins show unexpected localisation dynamics at centromeres (Dunleavy et al., 2012; Raychaudhuri et al., 2012). CAL1 is progressively reduced during prophase I and is undetectable for the remainder of meiosis. Contrastingly, centromeric CENP-C increases in prophase I and is undetectable in post-meiosis II spermatids, coincident with the second phase of CID assembly. Remarkably, CID is one of the few histones retained in the mature sperm of *Drosophila*, surviving histone-to-protamine exchange during spermiogenesis (Dunleavy et al., 2012; Raychaudhuri et al., 2012; Dubruille et al., 2025). Here, we investigate the significance of the unusual CAL1 and CENP-C dynamics during the first assembly phase in prophase I and determine the previously unexplored requirements for the second assembly phase on spermatids.

## Results

### Haploid spermatids have CID levels comparable to diploid spermatogonia

*Drosophila* testes provide a spatiotemporal array encompassing all stages of spermatogenesis, with germline stem cells (GSCs) located in the apex and later-developing stages towards the distal end (Figure 1A). *Drosophila* males undergo achiasmatic meiosis (Adams et al., 2020), lacking chromosomal features such as synapsis, homologous recombination and chiasmata formation (McKee et al., 1992). To account for these differences, alternative nomenclature has been introduced to distinguish stages of prophase I (Cenci et al., 1994). This convention subdivides prophase I into stages S1 to S6, characterised by an expansion in nuclear volume and the appearance of distinct chromosome territories into which homologous chromosome pairs are localised. Immediately after meiosis II, spermatids are described in stages T1 to T5, based on the appearance of a mitochondrial aggregate. As previously described (Dunleavy et al., 2012; Raychaudhuri et al., 2012), centromeric CAL1 progressively disappears during meiotic prophase I (Figure S1A, S1B), between S1 and S4, corresponding with a gradual increase in CID and CENP-C at centromeres (Figure 1A). After meiosis II, CID gradually increases during T1-T5 stages in spermatids, yet CENP-C progressively disappears and is absent at T5 (Figure 1A, S1C).

**Figure 1:**
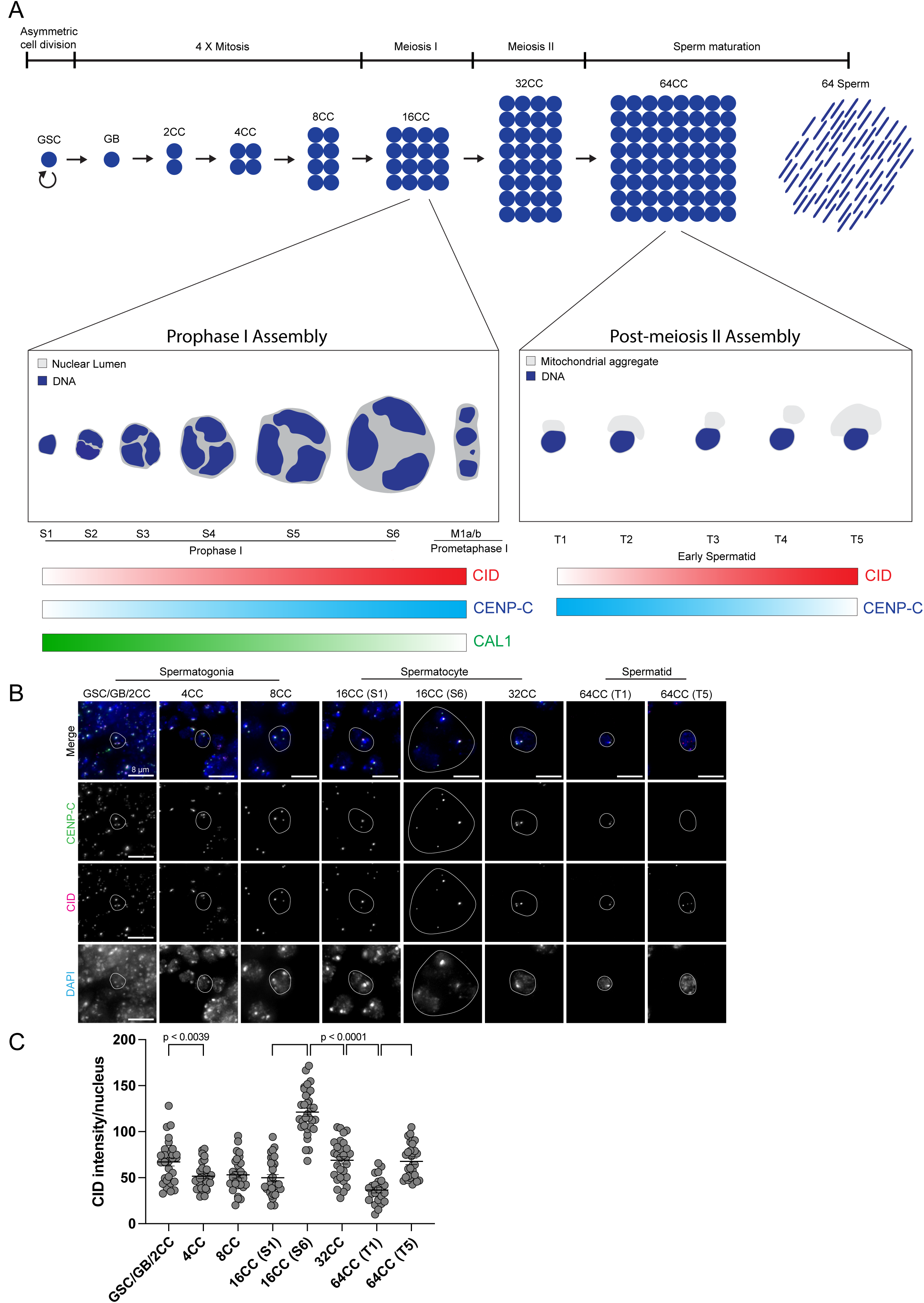
*Drosophila* melanogaster spermatogenesis and centromere assembly dynamics during meiosis. (**A**) Germline stem cells (GSC) divide asymmetrically generating a self-renewing GSC and a differentiating daughter cell called a gonialblast (GB). Four mitotic divisions with incomplete cytokinesis generates a cell cyst (CC) of 16 primary spermatocytes. Prophase I of meiosis is subdivided into six stages (S1 to S6), characterised by a massive expansion in nuclear volume, decondensation of DNA and the appearance of three chromosome territories (Cenci, 1994). CID and CENP-C are assembled onto the centromere during prophase I, coinciding with a loss of chaperone CAL1. Prometaphase I nuclei (stage M1a/b) are arranged as 3-4 condensed chromosome territories. Meiotic cells divide twice to separate homologous chromosomes and sister chromatids in meiosis I and II, respectively. CID is assembled again after meiosis II in stages T1 to T5 of the 64CC early spermatids, during which mitochondrial material is also removed from the cells. CENP-C is progressively lost with new CID deposition and is undetectable by stage T5. The resulting bundle of 64 early spermatids undergo drastic chromatin remodelling and individualise to form needle-shaped mature sperm nuclei. (**B**) *Drosophila* wild-type pupal testes stained with anti-CENP-C (green), anti-CID (magenta), and DAPI (blue). Representative images of cell stages during spermatogenesis from the 2CC spermatogonia to T5 stage early spermatids (64CC). A representative nucleus has been outlined in white for each stage. Cells were staged based on nuclear appearance and relative distance from the testis apex. S1 and S6 show early and late prophase I respectively. Scale bars = 8 µm. (**C**) Quantitation showing the relative fluorescent intensity per nucleus of the CID during each stage of spermatogenesis, with respective p-values annotated on the graph. n = 30 - 40. Error bars = SEM.

To assess overall CID dynamics across the germline, centromeric CID intensity was measured in spermatogonial 2-cell cysts (2CC, including GSC and gonialblast pairs), 4-cell cysts (4CC), 8-cell cysts (8CC), spermatocytes at early prophase I (16CC at S1) and at late prophase I (16CC at S6), 32-cell cysts that have completed meiosis I (32CC) and 64-cell cysts that have completed meiosis II (64CC at T1 and T5) (Figure 1A, 1B, 1C). A decrease in CID was measured between 2CC and 4CC, consistent with previous reports that GSCs harbour approximately 1.5-fold more CID than daughter cells (Ranjan et al., 2019; Kochendoerfer et al., 2023; Tower et al., 2025). CID levels were then quantitatively maintained throughout the subsequent mitotic divisions in 4CC, 8CC, and upon entry into meiosis in 16CC at S1 stage. Between S1 and S6 stages of prophase I, an approximate 2.5-fold increase in CID was noted, in line with the first phase of CID assembly occurring at this stage (Dunleavy et al., 2012; Raychaudhuri et al., 2012). After the first meiotic division, CID levels were reduced by approximately 50% in 32CC. This is expected given homologous chromosome segregation in meiosis I and a reduction in ploidy. A further proportional reduction was measured in 64CC at the T1 spermatid stage, congruous with sister chromatid segregation. The T1 stage of spermatid development marked the lowest levels of CID measured throughout spermatogenesis. Consistent with previous findings of a second phase of CID assembly in post-meiotic spermatids (Dunleavy et al., 2012), an increase in CID intensity (2.5-fold) was observed between T1 and T5 stages. Overall, CID levels in haploid T5 spermatids were equivalent to the levels in diploid 2CC stage spermatogonia. This result indicates that despite two rounds of chromosome segregation in meiosis, CID level in haploid spermatids is comparable to diploid pre-meiotic cells due to two unique phases of CID assembly occurring during spermatogenesis, with each providing higher than expected quantities (2.5-fold measured increases compared to the expected 2-fold).

### CENP-C localisation in meiosis is uncoupled from CID and CAL1 levels

In mitosis, CID, CAL1, and CENP-C are interdependent for centromere localization (Erhardt et al., 2008; Roure et al., 2019). To examine interdependencies in meiosis, the GAL4-UAS system was used to target CID, CAL1 or CENP-C mRNA for degradation, thereby reducing the pool of proteins available for deposition at centromeres. RNAi was induced using the germline-specific driver *bam*-GAL4 (active in 8CC spermatogonia and 16CC spermatocytes), and centromeres were stained with anti-CID, anti-CENP-C or anti-CAL1 antibodies. CID intensity was first quantified at the S1 stage, representing the start of CID assembly (Figure 1C). Quantitation of total centromeric CID intensity per nucleus showed normal levels in CID and CENP-C RNAi, confirming that CID assembly is not yet impacted (Figure 2A, 2B). CID and CENP-C RNAi displayed normal CENP-C levels, corresponding with normal levels of CID (Figure 2A, 2C). Contrastingly, centromeric CID was significantly reduced after CAL1 RNAi, indicating that CAL1 is required to stabilise CID already incorporated at centromeres at the S1 stage (Figure 2A, 2B). Despite this loss of CID, CENP-C levels were quantified as normal at S1 in the CAL1 RNAi (Figure 2A, 2C). This indicates that CID is not required to proportionally maintain the centromeric association of CENP-C in early prophase I. In the CAL1 RNAi, CAL1 was completely depleted from centromeres at S1 (Figure 2A, 2B), likely reflecting its higher turnover rate and lack of stable integration at the centromere (Mellone et al., 2011; Roure et al., 2019). Surprisingly, CAL1 was almost undetectable at the centromere after CID RNAi, demonstrating that CAL1 requires a newly synthesised CID for its localisation to centromeres at S1, where it functions to stabilise nucleosomal CID (Figure 2D, 2E). Contrastingly, in the CENP-C RNAi, CAL1 localised normally to centromeres and in the nucleolus (Figure 2D, 2E). In summary, at the S1 stage, CAL1 is required for the stability of both nucleosomal and soluble CID, and CENP-C can localise to centromeres in a manner that is not strictly dependent on CID and CAL1 level.

**Figure 2:**
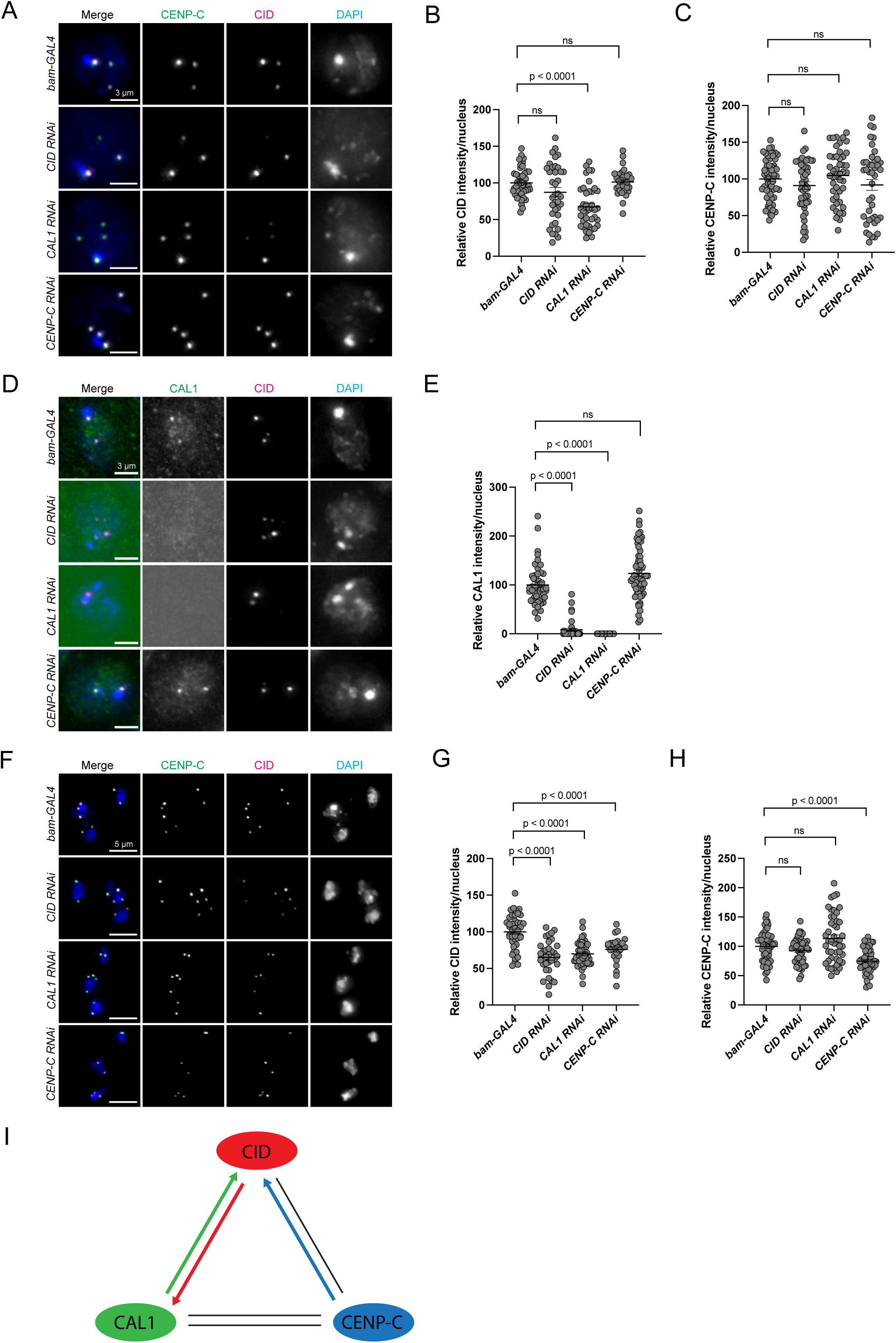
Requirements of CID, CAL1 and CENP-C for their recruitment and stability in meiosis I. (**A**) Pupal testes stained with anti-CENP-C (green), anti-CID (magenta) antibodies, and DAPI (blue). Representative images of early prophase I (S1 stage) cells from each of the genetic lines (*bam*-*GAL4, CID RNAi, CAL1 RNAi* and *CENP-C RNAi*). Scale bars = 3 µm. (**B**) Quantitation showing the relative fluorescent intensity per nucleus of CID at S1 in early prophase I. n = 46, 37, 39, 34. Error bars = SEM. (**C**) Quantitation showing the relative fluorescent intensity per nucleus of CENP-C at S1 in early prophase I. n = 57, 47, 30, 31. Error bars = SEM. (**D**) Early prophase I (S1 stage) cells stained with anti-CAL1 (green), anti-CID (magenta) antibodies, and DAPI (blue). Scale bars = 3 µm. (**E**) Quantitation showing the relative fluorescent intensity per nucleus of the histone chaperone CAL1 at S1 in early prophase I. n = 46, 37, 55, 61. Error bars = SEM. (**F**) Prometaphase I cells stained with anti-CENP-C (green), anti-CID (magenta) antibodies, and DAPI (blue). Each panel shows one nucleus, visible as 3 to 4 chromosome territories at prometaphase I. Scale bars = 5 µm. (**G**) Quantitation showing the relative fluorescent intensity per nucleus of CID at prometaphase I. n = 42, 33, 45, 27. Error bars = SEM. (**H**) Quantitation showing the relative fluorescent intensity per nucleus of CENP-C at prometaphase I. n = 50, 50, 46, 49. Error bars = SEM. (I) Model depicting the interdependencies for CID, CAL1 and CENP-C recruitment during meiosis I. Arrows represent a complete dependency between these proteins for their centromeric recruitment, with the colour indicating the direction of this dependency. Black lines indicate incomplete dependency.

We next examined CID intensity at prometaphase I, immediately after S6 and representing the maximal level of CID assembly (Figure 1C). Primary spermatocytes at prometaphase I were identified based on nuclear morphology, typically comprising 3-4 condensed chromosome territories (Figure 2F). Quantitation of total centromeric CID intensity per nucleus measured an approximate 35%, 30%, and 25% reduction in CID, CAL1, and CENP-C RNAi lines, respectively (Figure 2G), consistent with previous studies (Dunleavy et al., 2012). To offset CID dilution by half during DNA replication, new CID assembly should amount to 50% of the total pool in each cell cycle to maintain appropriate levels at the centromere. Our data indicate that only 35% of the total CID population at centromeres is replenished in prophase I, although it is possible that this underloading results from incomplete CID, CAL1, and CENP-C depletion in this time window. Next, centromeric CENP-C localisation was quantified per nucleus in CID, CAL1, and CENP-C RNAi to establish whether inner kinetochore recruitment was impacted. CENP-C was reduced by approximately 30% in the CENP-C RNAi, consistent with the 25% reduction in CID (Figure 2F, 2G, 2H). However, CENP-C level was unchanged in CID and CAL1 RNAi, despite previously observed reductions of 35% and 30% in centromeric CID (Figure 2F, 2G, 2H). This result indicates that despite a reduction in the total amount of CID at centromeres in all three RNAi lines, CENP-C assembly was affected only when it was targeted directly in the CENP-C RNAi. It also suggests that new CID assembly in prophase I is not required for CENP-C recruitment to meiotic centromeres. Furthermore, CID that is stably incorporated from prior mitotic divisions is sufficient to permit normal CENP-C recruitment in meiosis. Taken together, analyses of both early (S1) and late (prometaphase I) stages demonstrate that CENP-C localisation in meiosis is not strictly dependent on CID or CAL1. Thus, unlike mitosis, the interdependency feedback loop of centromere localisation between CID, CAL1, and CENP-C is incomplete (Figure 2I).

### CENP-C is more important for meiotic kinetochore recruitment and microtubule attachment than the underlying CID level

To determine the functional importance of CID assembly in prophase I, recruitment of the outer kinetochore protein Spindle Pole Component 105 (Spc105) was monitored in CID, CAL1, and CENP-C RNAi (Figure 3A, 3B). Spc105 is homologous to human Kinetochore Scaffold 1 (KNL1) and is required to maintain chromosome orientation and cohesion, recruit NDC80 and establish microtubule attachments (Joshi et al., 2024). Strikingly, Spc105 signal at centromeres in prometaphase I was greatly reduced only in the CENP-C RNAi, indicating that the assembly of new CENP-C in prophase I is necessary to recruit the outer kinetochore (Figure 3A, 3B). Thus, CENP-C inherited from prior divisions is not sufficient to establish intact meiotic kinetochores. Importantly, Spc105 recruitment was restored in a CENP-C rescue experiment in which CENP-C RNAi was performed in a line expressing RNAi-resistant HA-tagged CENP-C under UAS control (Figure 3C-3H). HA-CENP-C was confirmed to be expressed and incorporated at prometaphase I centromeres using anti-HA staining (Figure 3C). CID and CENP-C levels were also confirmed to be normal in the rescue flies, demonstrating that CENP-C deficiency drives the loss of Spc105 recruitment (Figure 3D, 3E, 3F). In contrast to the CENP-C RNAi, Spc105 was present at levels unexpectedly higher than the control in CID and CAL1 RNAi (Figure 3A, 3B). This result suggests that despite reduced CID at centromeres, CENP-C recruitment occurs in a manner that stimulates further Spc105 recruitment.

**Figure 3:**
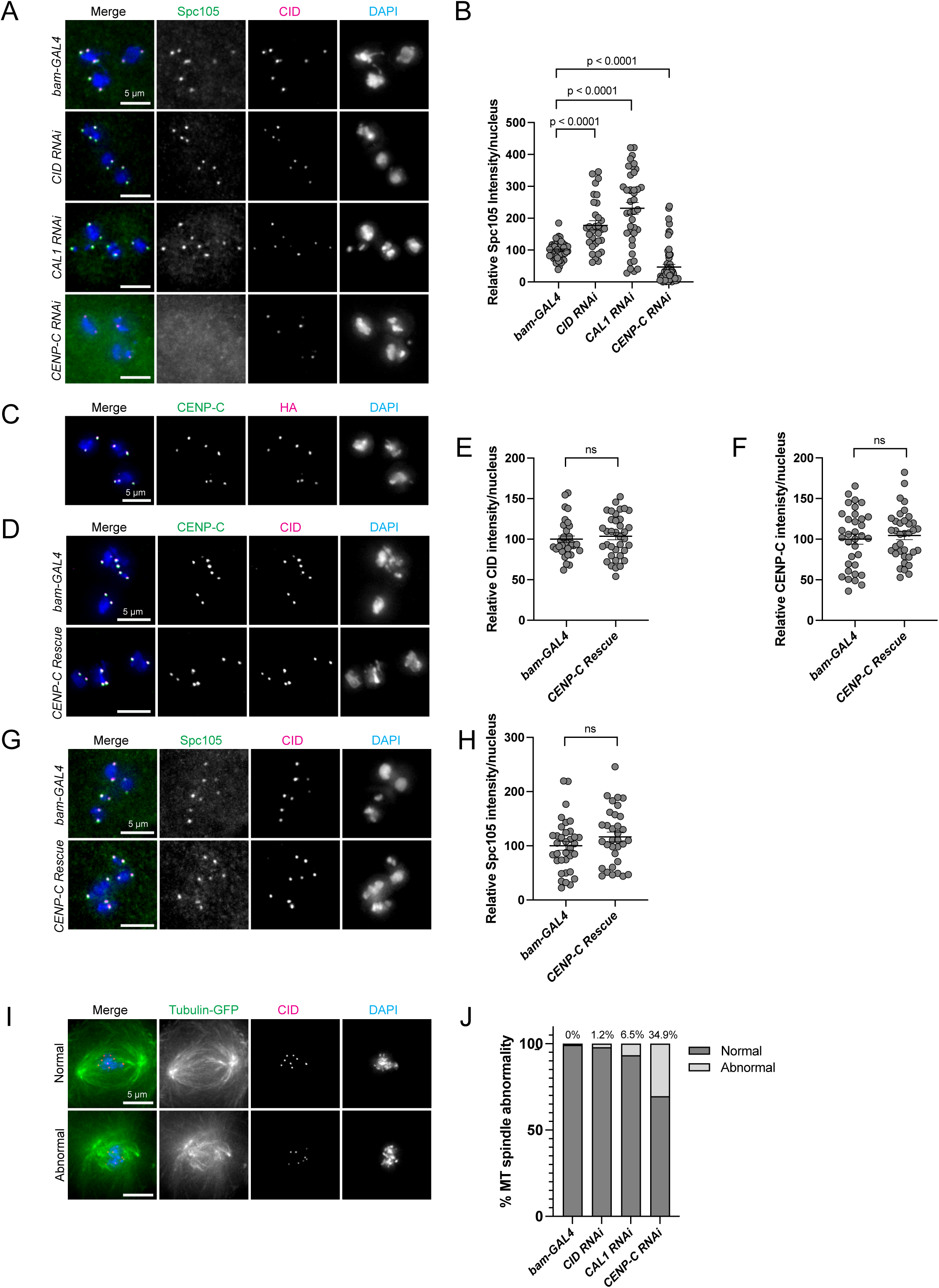
Outer kinetochore recruitment and microtubule spindle assembly after CID, CAL1 and CENP-C RNAi. (**A**) Pupal testes stained with anti-Spc105 (green), anti-CID (magenta) antibodies, and DAPI (blue). Representative images of prometaphase I cells from each of the genetic lines (*bam*-*GAL4, CID RNAi, CAL1 RNAi and CENP-C RNAi*). Scale bars = 5 µm. (**B**) Quantitation showing the relative fluorescent intensity per nucleus of the outer kinetochore component Spc105 (homolog of KNL1 in humans) at prometaphase I. n = 52, 32, 40, 57. Error bars = SEM. (C) Prometaphase I cell from CENP-C rescue flies (genotype = *bam*-*GAL4; UAS-CENP-C RNAi; UAS-HA-CENP-C*) stained with anti-CENP-C (green), anti-HA (magenta) and DAPI. Scale bar = 5 µm. (**D**) Representative images of prometaphase I cells from *bam*-*GAL4* and CENP-C rescue testes showing anti-CENP-C (green), anti-CID (magenta) and DAPI (blue). Scale bar = 5 µm. (**E**) Quantitation showing relative fluorescent intensity per nucleus of CID in *bam*-GAL4 control and CENP-C rescue cells at prometaphase I. n = 34, 36. (**F**) Quantitation showing relative fluorescent intensity per nucleus of CENP-C. n = 34, 36. (**G**) Representative images of prometaphase I cells from *bam-GAL4* and *CENP-C* rescue testes showing anti-Spc105 (green), anti-CID (magenta) and DAPI (blue). Scale bar = 5 µm. (**H**) Quantitation showing relative fluorescent intensity per nucleus of Spc105. n = 35, 33. (**I**) Representative images of normal and abnormal microtubule spindles showing tubulin-GFP (green), anti-CID (magenta) and DAPI (blue). Scale bar = 5 µm. (J) Quantitation of microtubule spindle abnormality in each knockdown line (*bam*-*GAL4 ; Tubulin-GFP/WT, CID RNAi/Tubulin-GFP, CAL1 RNAi and CENP-C RNAi*) with meiosis I and II pooled together.

Assembly of bipolar meiotic spindles was next monitored in the CID, CAL1, and CENP-C RNAi lines expressing GFP-tagged tubulin and immunostained for CID (Figure 3I, 3J). Spindles lacking a defined bipolar structure and without direct contact to centromeres were scored as abnormal for both meiotic divisions. Quantitation of normal and abnormal meiotic spindles revealed that 34.9% of meiotic figures were abnormal in the CENP-C RNAi, compared to 1.2% and 6.5% in the CID and CAL1 RNAi, respectively. Though CID and CAL1 RNAi cells have reduced centromeric CID, an apparent compensatory recruitment of CENP-C (and Spc105) prevented spindle abnormalities. Taken together, the assembly of meiotic CENP-C appears to dictate the level of kinetochore recruitment, independent of the underlying CID level. Additionally, a deficiency in CID and CAL1 can be tolerated, providing that sufficient CENP-C is recruited.

### CENP-C is crucial for accurate meiotic chromosome segregation

To investigate the consequences of centromere deficiencies on meiotic chromosome segregation, missegregation rates were examined in CID, CAL1, and CENP-C RNAi lines. To track segregation of homologous sex chromosomes in meiosis I, FISH probes recognising sequences specific to the X (AATAC repeat) and Y (359 bp repeat) chromosomes were used (Figure 4A, 4B). In primary spermatocytes, the Y chromosome probe typically displays two foci, whereas the X chromosome is visualised as a single focus, consistent with (Tsai et al., 2011). Pairs of secondary spermatocytes that just completed homologous sex chromosome segregation in meiosis I were scored as normal if one nucleus had an X chromosome and the other a Y chromosome. Any variation to this pattern was scored as abnormal. Pairs of spermatids that just completed meiosis II were scored as normal if both nuclei had an X chromosome, or both nuclei had a Y chromosome. Again, any variation to this pattern was scored as abnormal for meiosis II. To track segregation of autosomes in meiosis I and II, we used FISH probes specifically recognising sequences on the 2^nd^ (Responder, Rsp), 3^rd^ (Dodeca) or 4^th^ (AATAT repeat) chromosomes (Figure 4A, 4C). Pairs of spermatocytes that completed meiosis I or II were scored as normal if each nucleus had one of each focus, representing the segregation of homologous chromosomes and sister chromatids, respectively. Any variation to this pattern in meiosis I or II was scored as abnormal. For the sex chromosomes (Figure 4D), the highest chromosome missegregation rate was measured for CENP-C RNAi, in both meiosis I (28.6%) and II (49.4%). For the autosomes (Figure 4E), the highest chromosome missegregation rate was again measured for CENP-C RNAi, in both meiosis I (63.5%) and II (77.9%). Overall, CENP-C RNAi showed a chromosome loss rate of approximately 90%, with just 10% of post-meiotic spermatids displaying a normal karyotype (Figure S2). Elevated rates of autosome missegregation were counted in the CID RNAi (6.5% for meiosis I and 17.3% for meiosis II) and were increased in the CAL1 RNAi (13.7% for meiosis I and 19.9% for meiosis II). Thus, CID and CAL1 RNAi displayed missegregation rates higher than the control, but consistently lower than CENP-C RNAi. These results demonstrate that CENP-C levels in meiosis are crucial for completing correct chromosome segregation. Finally, the impact of CID, CAL1, and CENP-C depletion during meiosis on male fertility was tested. Despite observed deficiencies in centromeric CID, individualised mature sperm were visible at levels comparable to the control in the seminal vesicles for all three RNAi lines (Figure 4F). Fertility tests did not measure any defect in the number of adult progenies produced compared to the *bam*-GAL4 control (Figure 4G). In summary, despite reduced meiotic CID levels and high rates of meiotic chromosome missegregation, particularly in the CENP-C RNAi, CID, CAL1, and CENP-C depletion did not impact male fertility.

**Figure 4.**
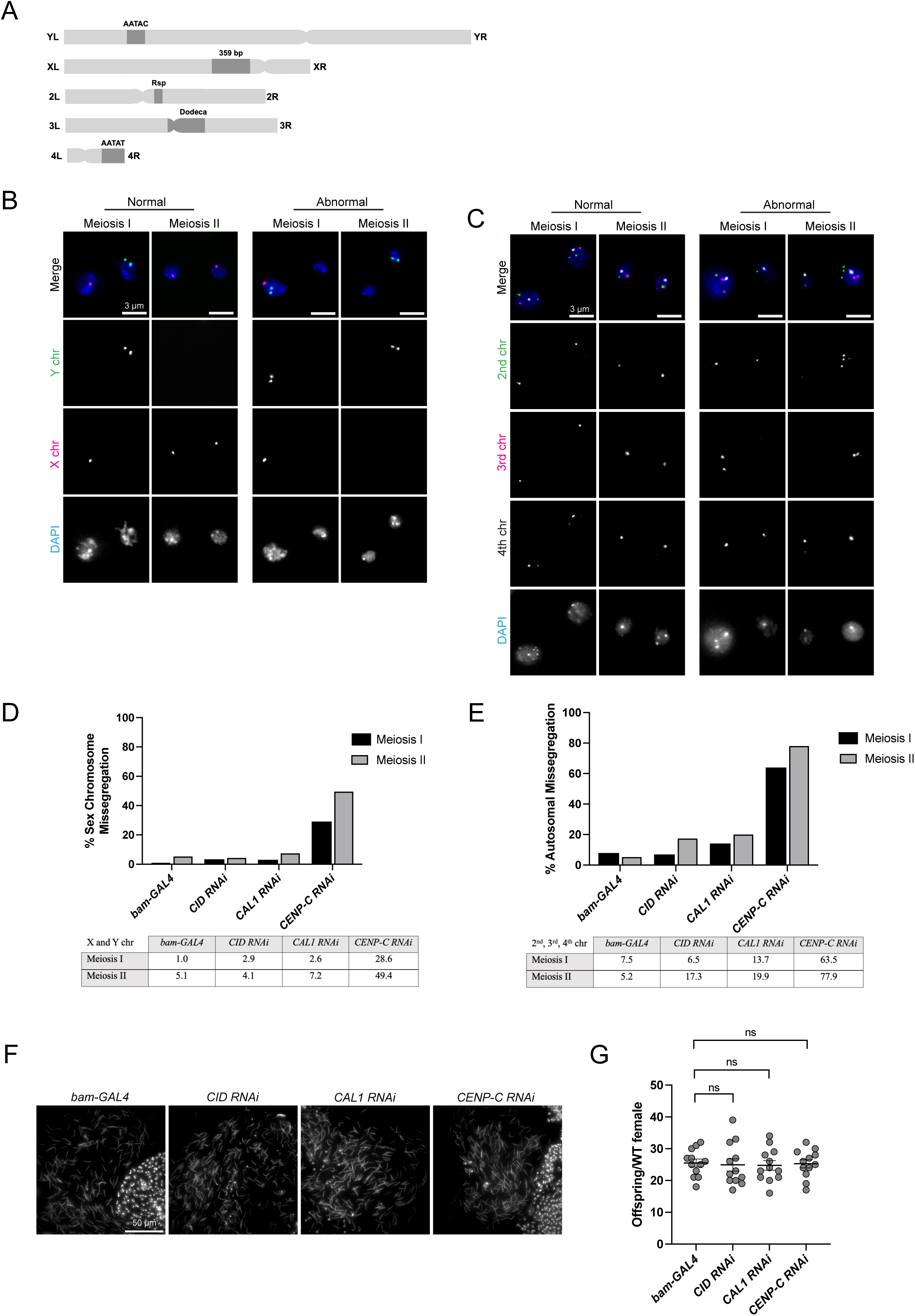
Meiotic chromosome missegregation and overall fertility after CID, CAL1 and CENP-C RNAi. (**A**) Schematic of the fluorescence *in situ* hybridisation probes used to follow chromosome segregation through the meiotic divisions. Probes are specific for each of the three autosomes and the sex chromosome pair (X and Y) in *Drosophila* melanogaster. (**B**) Fluorescence in situ hybridisation with Y chromosome (green) and X chromosome (magenta) probes, performed on pupal testes from the control and three knockdown lines (*bam*-*GAL4, CID RNAi, CAL1 RNAi* and *CENP-C RNAi*). Representative images of the normal and abnormal chromosome segregation patterns using the X and Y probes in meiosis I and II. Scale bar = 3 µm. (**C**) Fluorescence in situ hybridisation with 2^nd^ (Rsp probe), 3^rd^ (Dodeca satellite) and 4^th^ (AATAT repeat) autosomes, showing normal and abnormal segregation patterns in meiosis I and II. Scale bar = 3 µm. (**D**) Quantitation displaying the percentage of sex chromosome missegregation for the control and each of the knockdown lines (*bam*-*GAL4, CID RNAi, CAL1 RNAi and CENP-C RNA*i) after meiosis I and II. n = 4470. (**E**) Quantitation of the percentage of autosomal missegregation for each genetic line after meiosis I and II. n = 3830. (F) Representative images displaying mature sperm nuclei stained with DAPI (blue) that have spilt out of the seminal vesicles of 3-5 day old males from each genetic line (*bam*-*GAL4, CID RNAi, CAL1 RNAi* and *CENP-C RNA*i). Scale bar = 50 µm. (**G**) Results of fertility assays in which males of each genetic line have been crossed to wild-type (Oregon R) females. The offspring per female was counted from three independent replicates.

### The second post-meiotic phase of CID assembly cannot recover insufficiencies in the first phase

To determine whether meiotic centromere deficiencies could be replenished during the second assembly event, CID levels in T5 spermatids were quantified after CID, CAL1, and CENP-C RNAi driven by *bam*-GAL4 (Figure 5A, 5B). Transcription of most genes ceases in late prophase I of spermatogenesis, with few known exceptions (Vibranovski et al., 2010). Analysis of testis-specific single-cell RNA-sequencing data from the Fly Atlas (Li et al., 2022) project confirms that *bam*, *cid*, *cal1*, and *cenp-c* are downregulated after late prophase I (16CC at S6) and remain so during spermatid differentiation (64CC) (Figure S3A-E). This suggests that cells are unable to transcribe new RNA to recover from meiotic deficiencies in the second phase of centromere assembly, although some protein translation from pre-existing transcripts is possible. Quantitation of centromeric CID in T5 spermatids confirmed that levels remain reduced (Figure 5A, 5B). CID levels were reduced by 53%, 35%, and 44% in the CID, CAL1, and CENP-C RNAi lines, respectively, and CENP-C was undetectable. This result indicates that the depletion of CID, CAL1, and CENP-C using the *bam*-GAL4 driver results in reduced CID at centromeres at prometaphase I, and that CID remains reduced in post-meiotic T5 spermatids. Interestingly, both CID and CENP-C RNAi showed a further reduction in CID from the levels previously established in prometaphase I (Figure 2G), whereas CAL1 RNAi did not. CID RNAi exhibited a further decrease in CID from 65% of control levels at prometaphase I to 47% in T5 early spermatids. CENP-C RNAi displayed the largest difference, reducing from 75% of control levels at prometaphase I to 56% after meiosis II. However, a very minimal reduction was observed in the CAL1 RNAi by comparison, from 70% of the prometaphase I control to 65% of the T5 stage control levels suggesting it may not function during this second phase of assembly.

**Figure 5:**
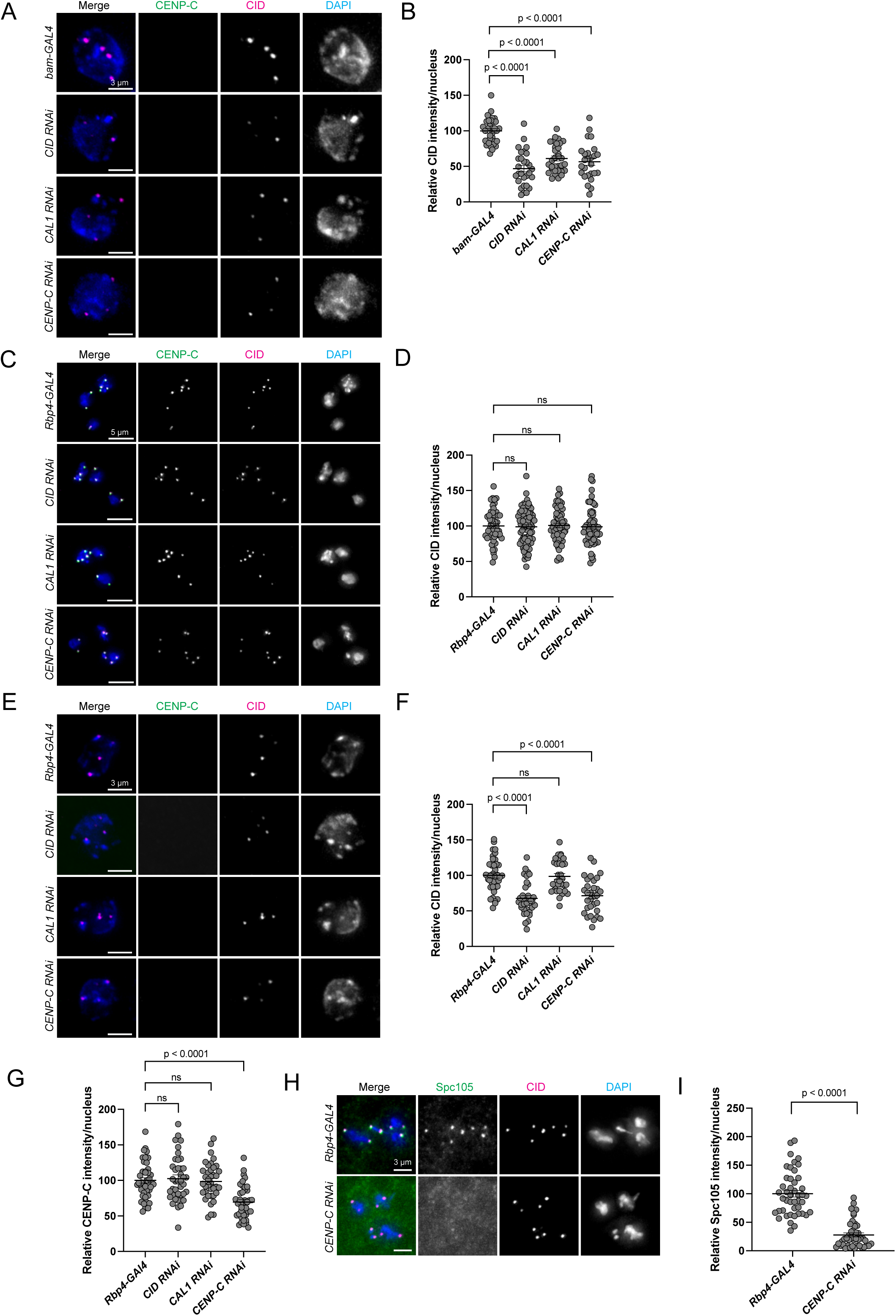
Requirements for the second phase of CID assembly after meiosis II. (**A**) Pupal testes stained with anti-CENP-C (green), anti-CID (magenta) antibodies, and DAPI (blue). Representative images of T5 spermatid nuclei from each of the genetic lines (*bam*-*GAL4, CID RNAi, CAL1 RNAi* and *CENP-C RNAi*). CENP-C staining is notably absent from each cell, verifying T5 stage spermatids identity. Scale bars = 3 µm. (**B**) Quantitation showing the relative fluorescent intensity per nucleus of CID. n = 32, 29, 27, 26. Error bars = SEM. (**C**) Pupal testes from a new GAL4 driver *Rbp4-GAL4* (expressed between S1 and S6) crossed to each RNAi line. Representative images of prometaphase I cells stained with anti-CENP-C (green), anti-CID (magenta) antibodies, and DAPI (blue) from each of the genetic lines (*Rbp4-GAL4*, *CID RNAi, CAL1 RNAi*, and *CENP-C RNAi*). Scale bars = 5 µm. (**D**) Quantitation showing the relative fluorescent intensity of CID per nucleus at prometaphase I. n = 50, 76, 57, 67. Error bars = SEM. (**E**) Representative images of early spermatid nuclei (T5 stage) stained with anti-CENP-C (green), anti-CID (magenta), and DAPI (blue) for each knockdown line and control (*Rbp4-GAL4*, *CID RNAi, CAL1 RNAi*, and *CENP-C RNAi*). CENP-C staining is absent, indicative of T5 stage spermatids. Scale bars = 3 µm. (F) Quantitation showing relative fluorescent intensity of CID per T5 nucleus. n = 39, 33, 32, 31. Error bars = SEM. (**G**) Quantitation showing the relative fluorescent intensity of CENP-C per nucleus at prometaphase I for each knockdown line and control (*Rbp4-GAL4*, *CID RNAi, CAL1 RNAi*, and *CENP-C RNAi*). n = 44, 39, 38, 39. Error bars = SEM. (H) Representative images of prometaphase I cells stained with anti-Spc105 (green), anti-CID (magenta) antibodies and DAPI (blue) from the genetic lines *Rbp4-GAL4* (control) and *Rbp4-GAL4*-induced CENP-C RNAi. (**I**) Quantitation showing the relative fluorescent intensity of Spc105 per nucleus at prometaphase I. n = 45, 44. Error bars = SEM.

### CID protein synthesis and assembly are uncoupled during spermatogenesis

To gain increased temporal resolution for RNAi depletion studies in prophase I, a second GAL4 driver was adopted. *Rbp4-GAL4* expression begins at the S1 stage in 16CC spermatocytes (Figure S3E), providing a narrower window for RNAi induction (Butsch et al., 2023). The functionality of this driver was first tested by driving expression of mCherry-tagged CID under UAS control. As expected, CID-mCherry foci were first detectable at the S1-2 stage of early prophase I, notably later than *bam*-GAL4 driven expression which appeared visible at 8CC (Figure S4A). At prometaphase I, CID-mCherry foci were visualised at centromeres, indicating that CID can be synthesised and assembled de novo at centromeres during prophase I (Figure S4B). Surprisingly, following CID, CAL1, and CENP-C RNAi using the *Rpb4-GAL4* driver, quantitation of centromeric CID at prometaphase I did not reveal any change in CID intensity (Figure 5C, 5D). This result differs from the approximate 30% reduction in CID that was measured in RNAi experiments using the *bam*-*GAL4* driver (Figure 2G). Thus, it appears that to significantly reduce levels of CID at the centromere during meiosis, RNAi constructs must be expressed in the prior cell cycle (8CC). Next, CID levels were quantified in T5 stage spermatids following *Rbp4-GAL4*-driven RNAi. Here, CID levels were reduced in both CID and CENP-C RNAi (Figure 5E, 5F), indicating that the CID protein translated in prophase I is assembled post-meiosis II. Interestingly, no significant change in CID was detected in T5 spermatids in the CAL1 RNAi, suggesting it does not impact the second phase of assembly (Figure 5E, 5F). Taken together, we conclude that CID protein synthesised during the 8-cell stage is loaded in prophase I (*bam*-*GAL4* RNAi result), while CID protein synthesised during the 16-cell stage is loaded following meiosis II (*Rbp4-GAL4* RNAi result).

### Functionally distinct pools of CENP-C exist in prophase I to support CID assembly and kinetochore recruitment

To assess requirements for inner kinetochore recruitment specifically in prophase I, CENP-C levels after CID, CAL1, and CENP-C RNAi using the *Rbp4-GAL4* driver were quantified. As with centromeric CID levels (Figure 5D), *Rbp4-GAL4*-driven CID and CAL1 RNAi did not significantly lower CENP-C levels at prometaphase I (Figure 5G). This resembles observations made with the *bam*-GAL4 driver (Figure 2F, 2H) and further uncouples CID, CAL1, and CENP-C recruitment interdependencies in meiosis. *Rbp4-GAL4*-driven CENP-C RNAi did display reduced CENP-C levels (Figure 5G), confirming that the *Rbp4-GAL4* driver and RNAi pathway were active in this time window and capable of reducing available proteins for deposition during prophase I. Thus, improved temporal resolution provided by the *Rbp4-GAL4* driver enabled the separation of CENP-C function between early and late prophase I. Early in prophase I, CENP-C is susceptible to *bam*-GAL4-driven depletion and appears to function primarily in CID assembly (Figure 2F, 2G). Contrastingly, later in prophase I, CENP-C is susceptible to both *bam*-GAL4- and *Rbp4-GAL4*-driven RNAi and switches its function towards recruitment of kinetochore proteins. As a demonstration of this, *Rbp4-GAL4*-driven CENP-C RNAi caused a reduction in Spc105 recruitment, despite normal CID levels (Figure 5H, 5I). This provides further evidence that CENP-C recruitment dictates outer kinetochore assembly in late prophase I, separate from underlying CID.

### CAL1-independent CID assembly after meiosis II requires CENP-C

Unlike CID or CENP-C RNAi driven by *bam*-GAL4, CAL1 RNAi did not cause further reductions in CID levels at T5 after the second phase of assembly (Figure 5A, 5B). Together with the absence of CAL1 from centromeres beyond prophase I, this result raised questions regarding the role of CAL1 during the post-meiotic CID loading event. Centromeric CID levels were not impacted at prometaphase I in *Rbp4-GAL4*-driven CAL1 RNAi (Figure 5C, 5D). In fact, CID levels remained indistinguishable from those of the control even after the second phase of CID assembly in the early T5 spermatids (Figure 5E, 5F). This indicates that CID is loaded in a manner independent of its chaperone CAL1 after meiosis II. CENP-C disappears from centromeres in the window during which CID is assembled after meiosis II (Figure S1C). To establish if CENP-C is required for this second phase of assembly, CID levels were quantified after *Rbp4-GAL4*-driven CENP-C RNAi. CENP-C levels were significantly reduced at prometaphase I, even as CID levels remained unchanged (Figure 5G). Moreover, CID was significantly reduced at centromeres in post-meiotic T5 stage spermatids, confirming that the pool of CENP-C synthesised later in prophase I is required for the second phase of CID assembly, as well as kinetochore formation at prometaphase I (Figure 5E, 5F). Taken together, these results demonstrate that CID assembly after meiosis II occurs independent of CAL1, but requires CENP-C.

## Discussion

Building on previous findings (Dunleavy et al., 2012; Raychaudhuri et al., 2012), we refine the requirements for centromere assembly in meiosis. We separate assembly events occurring in early and late prophase I, with and without CAL1. In early prophase I, CAL1 functions in CID assembly together with CENP-C. At late prophase I, CAL1 is no longer detectable, and CID assembly is complete. At this point, CENP-C continues to be recruited to the centromere, but it is no longer required for CID assembly. Rather, it functions in kinetochore recruitment. This pool of CENP-C also functions in the second phase of CID assembly, that uniquely, is independent of CAL1.

### Refining the requirements for CID assembly in prophase I

This study reveals a mutual dependency between CAL1 and CID in terms of stability and localisation in early prophase I. At S1 stage, CID depletion caused a dramatic reduction in CAL1, demonstrating that CAL1 requires newly synthesised non-nucleosomal CID for its stability and nuclear localisation. Reciprocally, following CAL1 RNAi, centromeric CID levels were significantly reduced at S1. This suggests that CAL1 further functions to stabilise nucleosomal CID already incorporated at the centromere at this time. Similar observations were reported in mitotic cells, where CENP-A-HJURP and CID-CAL1 pre-nucleosomal complexes offer reciprocal stabilisation in humans and flies, respectively (Erhardt et al., 2008). This is the first indication that similar mutual stabilisation dynamics occur in meiosis.

We also demonstrate that in contrast to mitosis (Erhardt et al., 2008), CENP-C recruitment in meiosis is not entirely dependent on CID or CAL1 levels. According to the current mitotic model, CAL1 complexed with CID is initially recruited to the centromere by pre-existing stably bound CENP-C. Newly localised CAL1 acts to recruit further CENP-C to the centromere, closing the epigenetic feedback loop (Roure et al., 2019). Here we find that at S1 stage, CENP-C can localise to centromeres in the absence of CAL1. Similarly, at prometaphase I, CENP-C can localise to centromeres at normal levels, despite lower levels of underlying CID. These results reveal that in meiosis the recruitment of CENP-C by CID nucleosomes is not stoichiometric. The mechanism by which additional CENP-C is recruited to the centromere in the absence of a portion of CID nucleosomes is not understood, but could derive from the dimerisation potential of CENP-C (Medina-Pritchard et al., 2020). Additionally, it is possible that under standard conditions not every CID nucleosome recruits one CENP-C molecule. This permits a saturation of CID with CENP-C only when certain CID nucleosomes are lost, providing a mechanism by which CENP-C can be recruited to a level that prevents meiotic chromosome segregation defects. Such a mechanism might explain why Spc105 recruitment was enhanced in the CAL1 and CID RNAi despite a 30% reduction in CID.

### CID assembly post-meiosis II is independent of CAL1, but requires CENP-C

This study establishes for the first time the requirements for the post-meiotic phase of CID assembly. CAL1 cannot be visualised at the centromere from mid-prophase I, remains undetectable for the rest of spermatogenesis and is absent from mature sperm (Raychaudhuri et al., 2012). Here we show that CAL1 depletion does not specifically impact CID levels in T5 spermatids, indicating that CID deposition occurs normally despite the absence of its chaperone. This represents the first known example of a CAL1-independent deposition event in flies, or indeed any species. It is interesting to speculate whether such an assembly could occur independently of HJURP in mammals. Furthermore, we show that this second phase of CID assembly is dependent on CENP-C. No evidence exists that CENP-C can assemble CID nucleosomes directly or that CID is capable of self-assembly, thus, we suggest that CENP-C is required to recruit unknown or canonical histone assembly factors for this purpose. The function of this second phase of deposition may be to prime spermatid cells for the drastic chromatin remodelling event involved with protamine exchange, during which CID remains one of few histones retained on the mature sperm (Raychaudhuri et al., 2012). Indeed, our analysis of the overall CID dynamics in the germline indicates that the level of CID on T5 spermatids is two-fold higher than expected for a haploid cell. Precisely how *Drosophila* CID is maintained on mature sperm in the absence of both CENP-C and CAL1 requires further investigation.

### Functionally distinct pools of meiotic CENP-C are required for CID assembly and kinetochore recruitment

We also identify distinct pools of CENP-C with separate, non-overlapping functions in meiosis. In early prophase, one CENP-C pool functions in CID assembly, presumably in the recruitment of CAL1. At prometaphase I, another CENP-C pool assembles the kinetochore. Furthermore, this pool of CENP-C does not function in CID assembly at that time. It is likely that CENP-C preferentially functions in kinetochore recruitment later in prophase I because CAL1 is absent. We show that CENP-C loading at prometaphase I is critical for Spc105 recruitment, which in turn is crucial for meiotic chromosome segregation. These findings parallel observations in *Drosophila* females in which a pool of CENP-C maintains centromeric CID in oocytes, which can be exchanged for a new and functionally distinct pool of CENP-C in prophase I (Fellmeth et al., 2023). Remarkably, in males, this ‘kinetochore’ pool of CENP-C takes on an additional function in post-meiosis II, where it is required for CID assembly on spermatids. This deposition event occurs in the absence of CAL1, and it is possible that CENP-C functions in the recruitment of a novel CID chaperone and assembly factor.

### Synthesis and deposition of CID protein are uncoupled during spermatogenesis

In mitosis, CID/CENP-A is primarily synthesised in G2-phase and deposited at the end of mitosis in the same cell cycle (Jansen et al., 2007; Schuh et al., 2007; Mellone et al., 2011). Here, we identify that the timing of CID synthesis and its cell cycle assembly are uncoupled during spermatogenesis. For the first phase of assembly in meiotic prophase I, most of the newly assembled CID protein appears to be synthesised in the preceding mitotic division. The temporal resolution of RNAi depletion experiments allowed us to conclude that the pool of CID assembled during prophase I is transcribed and translated at the 8CC stage. We note that if excess CID is provided in prophase I (via CID-mCherry overexpression driven by the *Rbp4-GAL4* driver), it can load at the centromere. However, under normal expression conditions, we propose that the CID pool synthesised at the 8CC stage is preferentially loaded. For the second phase of assembly in post-meiotic spermatids, we found that newly assembled CID protein is synthesised in meiotic prophase I. Exactly how soluble CID protein is chaperoned to survive both meiotic divisions is not known, but it is unlikely to work via CAL1, which is undetectable at this stage. This is an intriguing example of a protein synthesised in one cell cycle and used in the next, possibly facilitated by incomplete cytokinesis occurring in germline cysts. Comparable to most species, *Drosophila* spermatocytes undergo successive meiotic divisions with a very short intervening interkinesis (Fuller, 1998; Giansanti and Fuller, 2012). The lack of proper gap phases, during which CID protein is normally synthesised, may necessitate pre-translation of any protein required shortly after the second division.

### Conserved CID and CENP-C dynamics in the male and female germline

This work highlights a critical function for CENP-C recruitment in male meiosis. Strikingly, despite an approximate 50% reduction in CID in T5 spermatids and only 10% of spermatids displaying a normal karyotype in CENP-C RNAi, we did not measure any defect in male fertility (Figure 5G). This result is surprising, though it likely reflects the amplificatory nature of spermatogenesis; each GSC gives rise to a cyst of 64 spermatids, with multiple cysts in each testis. Therefore, despite high rates of meiotic chromosome missegregation, one normal sperm manages to fertilise the egg. To offset the 50% reduction in CID, it is possible that there is a compensatory loading of CID on paternal chromosomes in the embryo, which is proposed to occur in CENP-C depleted mouse oocytes (Tower et al., 2025). *Drosophila* male meiosis is atypical, lacking the signature features of meiosis of synapsis and recombination (Cenci et al., 1994) and therefore might not reflect centromere dynamics in meiosis in general. However, our results align well with the dynamics of CENP-C reported in female mice and *Drosophila*, which exhibit the classical features of meiosis (Smoak et al., 2016; Fellmeth et al., 2023). In female flies, it was proposed that replenishing the pool of CENP-C is necessary due to the extended arrest in prophase I that occurs in oocytes (Fellmeth et al., 2023). Yet, male meiosis does not encompass a prophase I arrest in any species. Therefore, mechanisms of CID assembly and CENP-C turnover are more likely to be related to a general switch in CENP-C function at prometaphase I that is critical for both sexes.

## Material and Methods

### Drosophila Stocks and husbandry

Fly stocks were propagated at 25°C on Nutri-Fly® standard medium supplemented with 0.5% propionic acid and 1% nipagin under a 12-hour light-dark cycle. These stocks were obtained from various sources as documented here. From Vienna *Drosophila* Research Centre, UAS-CID RNAi (VDRC #102090), *UAS-CAL1* RNAi (VDRC #45248), UAS-CENP-C RNAi (VDRC #33790, discontinued, but exists with the same construct ID 10208 as VDRC #33792). From Bloomington *Drosophila* Stock Centre, wild-type Oregon R (BDSC #25211), *bam*-GAL4 (BDSC #80579), *Rbp4-GAL4* (BDSC #600281). The fly line *bam*-GAL4; Tubulin-GFP was generated in the lab by a series of crosses between BDSC #80579 and Kyoto Stock Center #109603. From various other sources, the overexpression lines *UAS-CID*-mCherry (Dattoli et al., 2020), *UASp-HA-CENP-C/TM6* Tb (originally gifted as *UASp-HA-CENP-C/TM3* Sb from Kim McKim) and the CENP-C rescue line *UASp HA-CENP-C ; UAS-CENP-C* R*NAi* (Carty et al., 2021). To perform an experiment, virgin females from the specific GAL4 driver lines were mated with males displaying the appropriate markers from the RNAi or overexpression responder lines and the progeny were let develop at 29°C. F_1_ progeny were dissected at the early pupal stage of development to enrich for meiotic cells and early spermatid stages. Three biological replicates were performed for each experiment.

### Drosophila testis tissue preparation

Testes (adult and pupal) were dissected in 1X PBS and fixed on a SuperFrost slide in 4% paraformaldehyde (diluted with 1X PBS) for 10 minutes. All common reagents were obtained from Fisher Scientific or Merck unless otherwise specified. After fixation, the samples were gently squashed beneath a hydrophobic RainX-treated coverslip and snap-frozen in liquid nitrogen. The coverslip was quickly removed and the samples stored in 70% ethanol at -20°C until further processing. Processing for immunofluorescence and fluorescence *in situ* hybridisation involved passing through an ice-cold ethanol series (2 minutes each in 75%, 85% and 95% ethanol), allowing the samples to air dry for approximately 2 minutes and permeabilisation for 10 minutes in 1X PBS 0.4% Triton-X100 (PBST 0.4%) at room temperature.

### Immunofluorescence

Samples were blocked with 1% bovine serum albumin in 1X PBS for 1 hour at room temperature. Primary antibodies were diluted in blocking buffer and incubated with the samples overnight at 4°C. The next day, the samples were washed 3 x 10 minutes in 1X PBST 0.4% and incubated with secondary antibodies (1:500 dilution in blocking buffer) for 1 hour at room temperature in the dark. Then a further 3 x 10-minute washes in PBST 0.4% were performed. Samples were treated with DAPI (1ug/ml in 1X PBS) for 5 minutes at room temperature and washed in 1X PBS for 10 minutes. Finally, they were mounted in SlowFade^TM^ Gold antifade reagent (Invitrogen) and stored at -20°C.

### Antibodies

Primary antibodies used for immunofluorescence include: rat anti-CID (Active Motif #61735; used at 1:500), sheep anti-CENP-C (Dattoli et al., 2020; used at 1:500), sheep anti-CAL1 (Kochendoerfer et al., 2023; used at 1:1000), sheep anti-Spc105 (gifted from Marcin Przewloka, Venkei et al., 2011; used at 1:500), and mouse anti-HA (ThermoFisher #26183; 1:500). Species specific AlexaFluor 488, 546, 550 and 647 secondary antibodies were used at a standard 1:500 dilution.

### Fluorescence in situ hybridisation

To investigate homologous chromosome and sister chromatid segregation during meiosis I and II, fluorescently labelled probes for the X chromosome (359 bp repeat), Y chromosome (AATAC), 2^nd^ (Responder), 3^rd^ (Dodeca satellite) and 4^th^ chromosome (AATAT) were used.

Probes were synthesised as single-stranded oligonucleotides conjugated with 5’ ATTO fluorophores by Eurofins using sequences obtained from Tsai et al., 2011 and Ferree et al., 2009. Samples were washed for 3 x 10 minutes in 2X saline-sodium citrate-Tween 0.1% (2X SSC-Tw 0.1%) at room temperature, followed by one 10-minute wash in 2X SSC-Tw 0.1% + 25% formamide and 10 minutes in 2X SSC-Tw 0.1% + 50% formamide at room temperature. Pre-hybridisation was carried out in 2X SSC-Tw 0.1% + 50% formamide at 37°C for 2 hours. A quantity of 20 ng was used per sample for the 2^nd^, 3^rd^, X and Y chromosomes probes and40 ng was used for the 4^th^ chromosome probe. The probes were added to 1X hybridization buffer (3X SCC, 10 % dextran sulphate) with 50% formamide. 20 µl of hybridization buffer containing the respective FISH probes were added to each coverslip. Samples were inverted over the coverslips and sealed with Marabu Fixogum rubber cement. Denaturation was completed at 90°C for 4 minutes and followed with incubation overnight in a humidified chamber at 20°C. The coverslip and rubber cement were removed and a series of post-hybridisation washes were completed as follows: 1 x 10 minute wash at 20°C in 2X SSC-Tw + 50% formamide, 2 X 30 minute washes at 20°C in 2X SSC-Tw 0.1% + 50% formamide, 1 x 10 minute wash in 2X SSC-Tw 0.1% + 25% formamide at room temperature and 3 x 10 minute washes in 2X SSC-Tw 0.1% at room temperature. Samples were treated with DAPI for 5 minutes, washed in 1X PBS for 10 minutes, mounted in SlowFade^TM^ Gold antifade reagent and stored at -20°C.

### Microscopy and image analysis

Experiments were imaged by the Delta Vision Elite microscope (Applied Precision, Imsol) using primarily the 40X and 60X oil immersion Olympus X-line UPLXAPO40X 1.4NA and 1.42NA objectives. Z-stacks of 0.2 μm step size were used. Fluorescence passed through a 430–455 nm, 490–540 nm, 575–620 nm, 655–755 nm bandpass filter for detection for DAPI and Alexa Fluors 488, 546 and 647, respectively. Exposure times and transmission percentages were kept constant through each experiment. Images acquired at the microscope were deconvolved (conservative approach) using SoftWorx application (Applied Precision, Imsol) and processed in FIJI Image J.

### Image analysis

To quantify protein levels in the samples, fluorescent intensity was measured using FIJI Image J. Single nuclei were isolated amongst the images (8-bit) and Z-projected with maximal intensity, ensuring to capture the entirety of the centromeric signal. All images were saved as .tiff files. The background was subtracted (rolling ball radius set to 50.0) from the projected images and thresholding using the default algorithm was applied to select the entirety of the centromeric signal. Integrated density (mean gray value x area) was summed to generate the total fluorescent intensity per nucleus. Statistical analyses were performed using GraphPad Prism software. Each sample population was tested for normality, and corresponding parametric or non-parametric statistical tests were performed accordingly (Welch’s t-test or Mann-Whitney U test). P-values are displayed on each graph.

### Fertility assays

One virgin wild-type female was crossed to one RNAi male at 29°C and the total number of offspring per wild-type female was scored after 14 days.

## Acknowledgments

RSK is funded by Government of Ireland Postgraduate Scholarship GOIPG/2023/3987. EMD is funded by Research Ireland Frontiers for the Future Award 20/FFP-A/8519.

---

**Figure S1:**
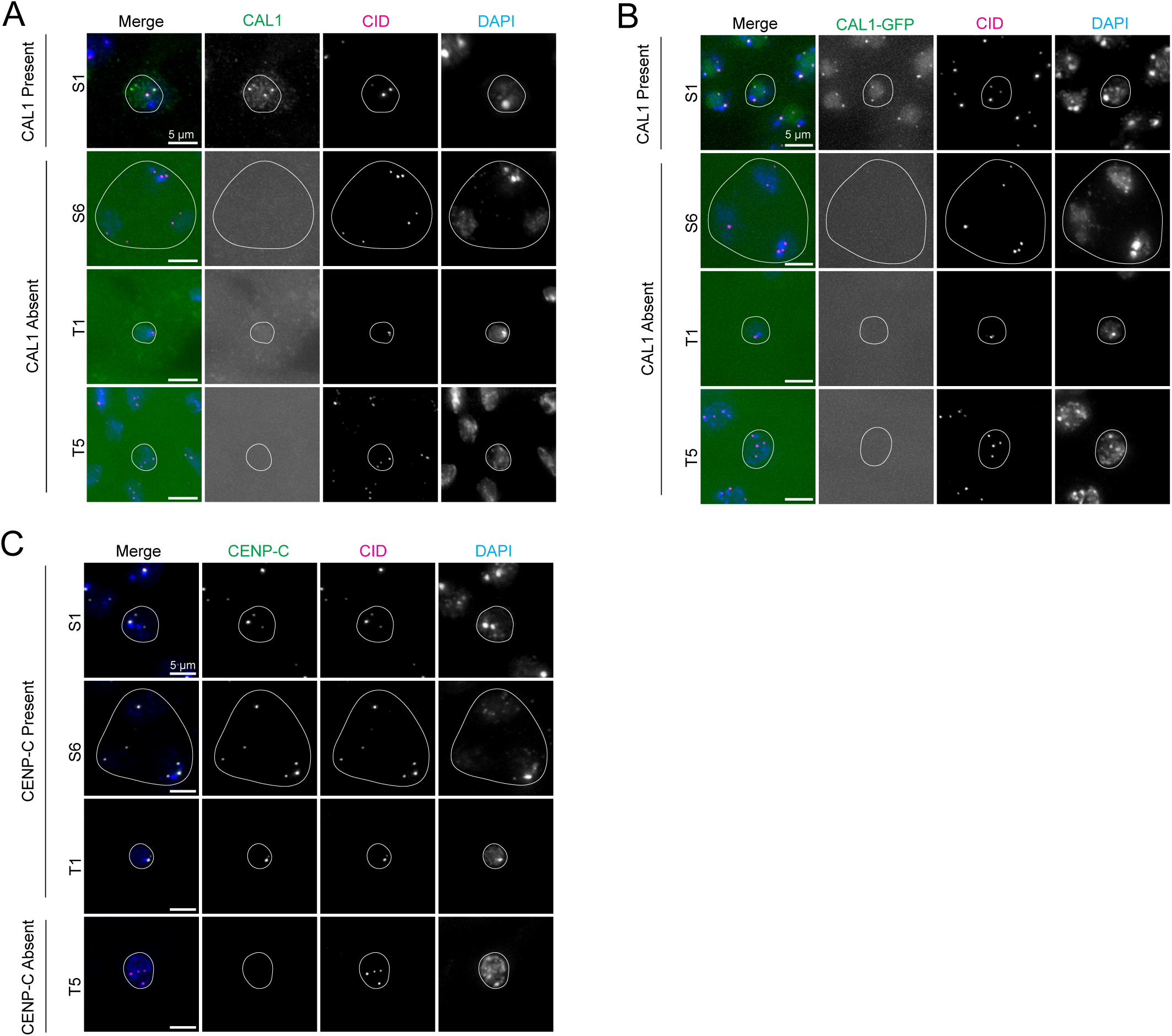
CAL1 and CENP-C recruitment dynamics during meiosis and in the early spermatids. (**A**) Pupal testes stained with anti-CAL1 (green), anti-CID (magenta) antibodies, and DAPI (blue). CAL1 is present at the centromeres of S1 stage cells (16CC) and absent from the centromeres of S6 (16CC), T1 and T5 (64CC). CAL1 is also visible in the nucleolus of S1 cells. Scale bars = 5 µm. (**B**) Pupal testes from CAL1-GFP flies stained with anti-CID antibody (magenta) and DAPI (blue), showing presence at S1 stage and in the nucleolus, as well as an absence from later stages (S6, T1 and T5). Scale bars = 5 µm. (**C**). Pupal testes stained with anti-CENP-C (green), anti-CID (magenta) antibodies, and DAPI (blue). CENP-C is present at the centromeres of S1, S6 and T1 stage cells, but absent from T5 stage centromeres. Scale bars = 5 µm.

**Figure S2:**
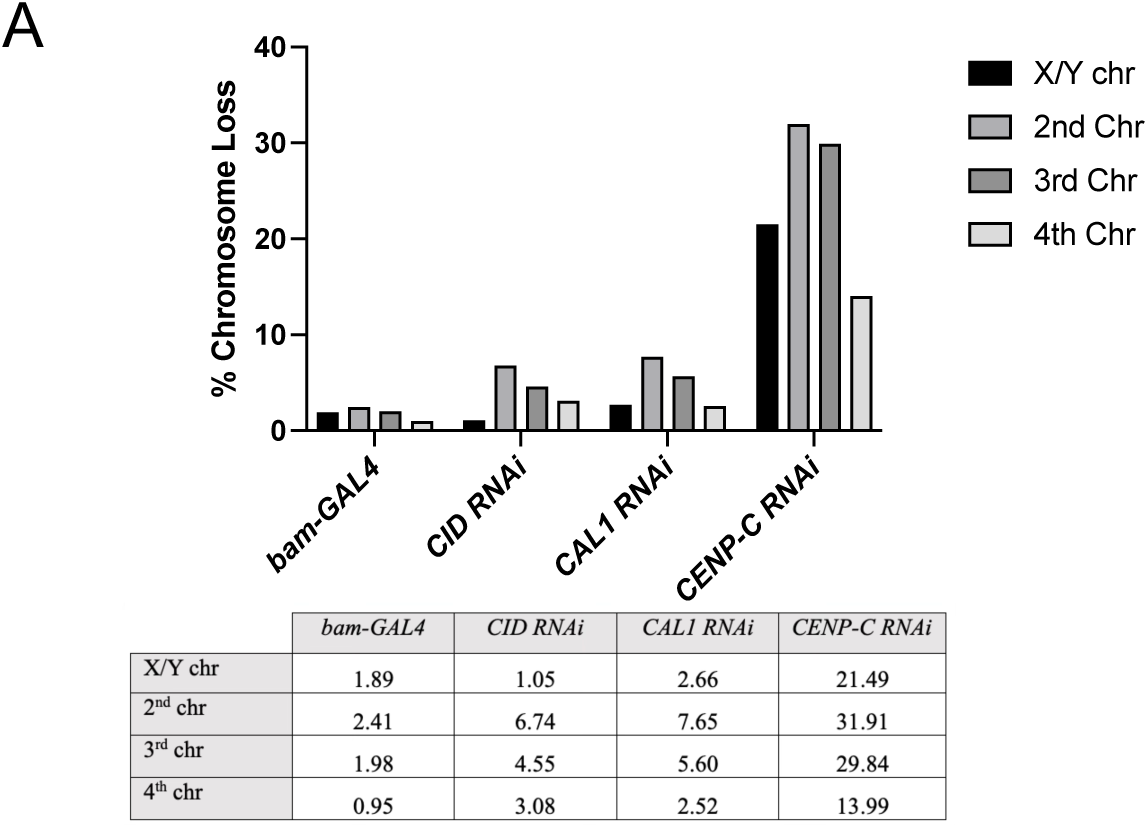
Chromosome loss upon CID, CAL1 and CENP-C RNAi. Quantitation of percentage chromosome loss for each homologous pair (X/Y, 2^nd^, 3^rd^ and 4^th^ chromosomes) in each knockdown line (*bam*-*GAL4, CID RNAi, CAL1 RNAi* and *CENP-C RNAi*). n = 8300.

**Figure S3:**
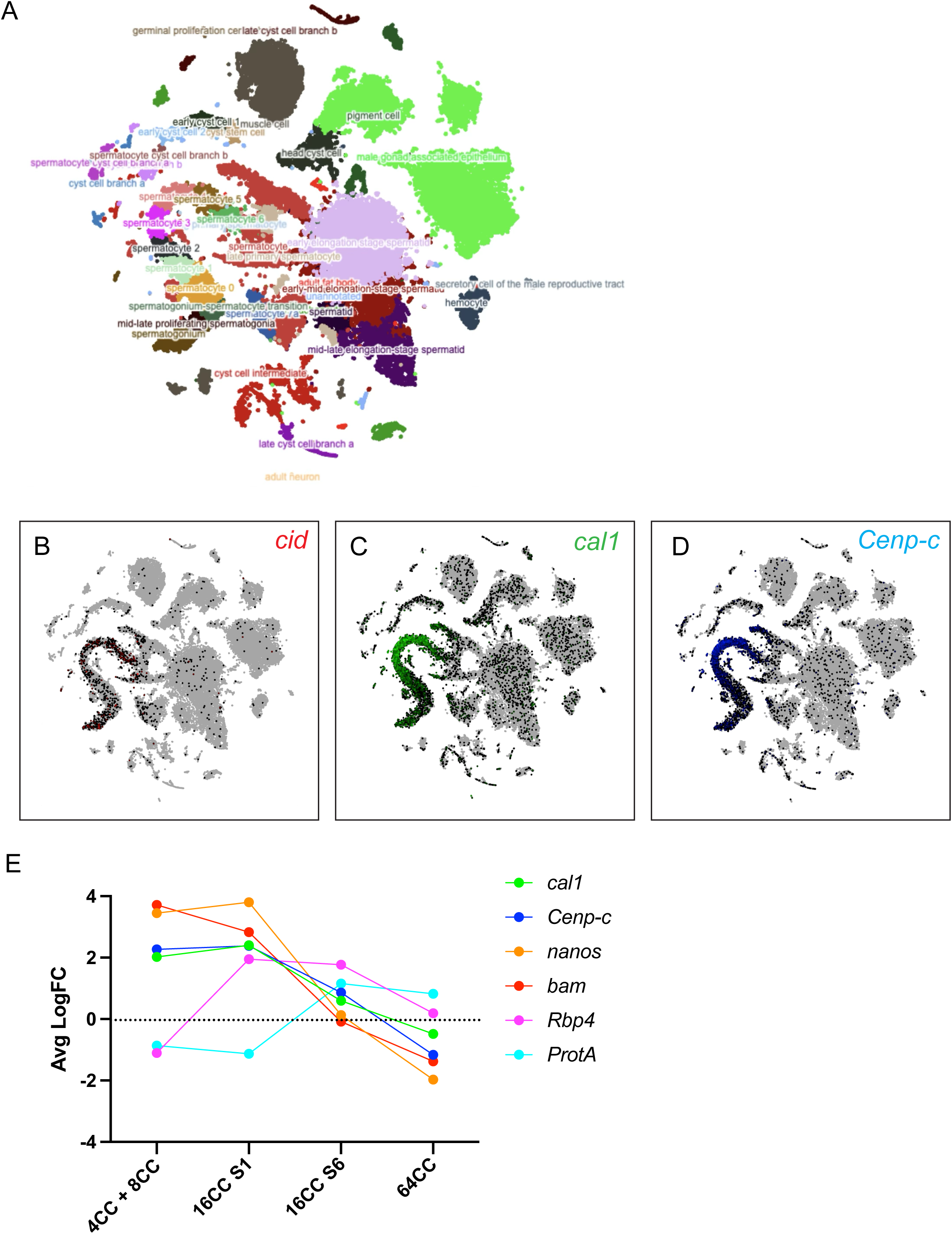
Single-cell RNA seq expression data for centromere proteins and known markers of spermatogenesis. (**A**) Uniform Manifold Approximation and Project (UMAP) graph displaying annotated testis cell clusters from the Fly Cell Atlas (Li et al., 2022), visualised using the SCope platform. (**B**) UMAP graph indicating expression levels of cid in adult testis. Black represents cells with lower expression and red shows higher expression. (**C**) UMAP graph displaying cal1 expression within the adult testis in green. (**D**) cenp-c expression within the adult testis shown in blue. (E) Graph depicting the average Log fold-change of cal1 and cenp-c, along with several known markers of spermatogenesis including nanos and *bam* (spermatogonia and spermatocytes), Rbp4 (spermatocytes), Protamine A (spermatocytes and spermatids). 4CC + 8CC data were derived from the mid-late proliferating spermatogonia cluster in the 10X dataset. Spermatocyte 1 and 6 clusters were mapped to early and late prophase I using known expression patterns of nanos, *bam* and Rbp4. Early-mid elongation stage spermatids were used as the 64CC dataset.

**Figure S4:**
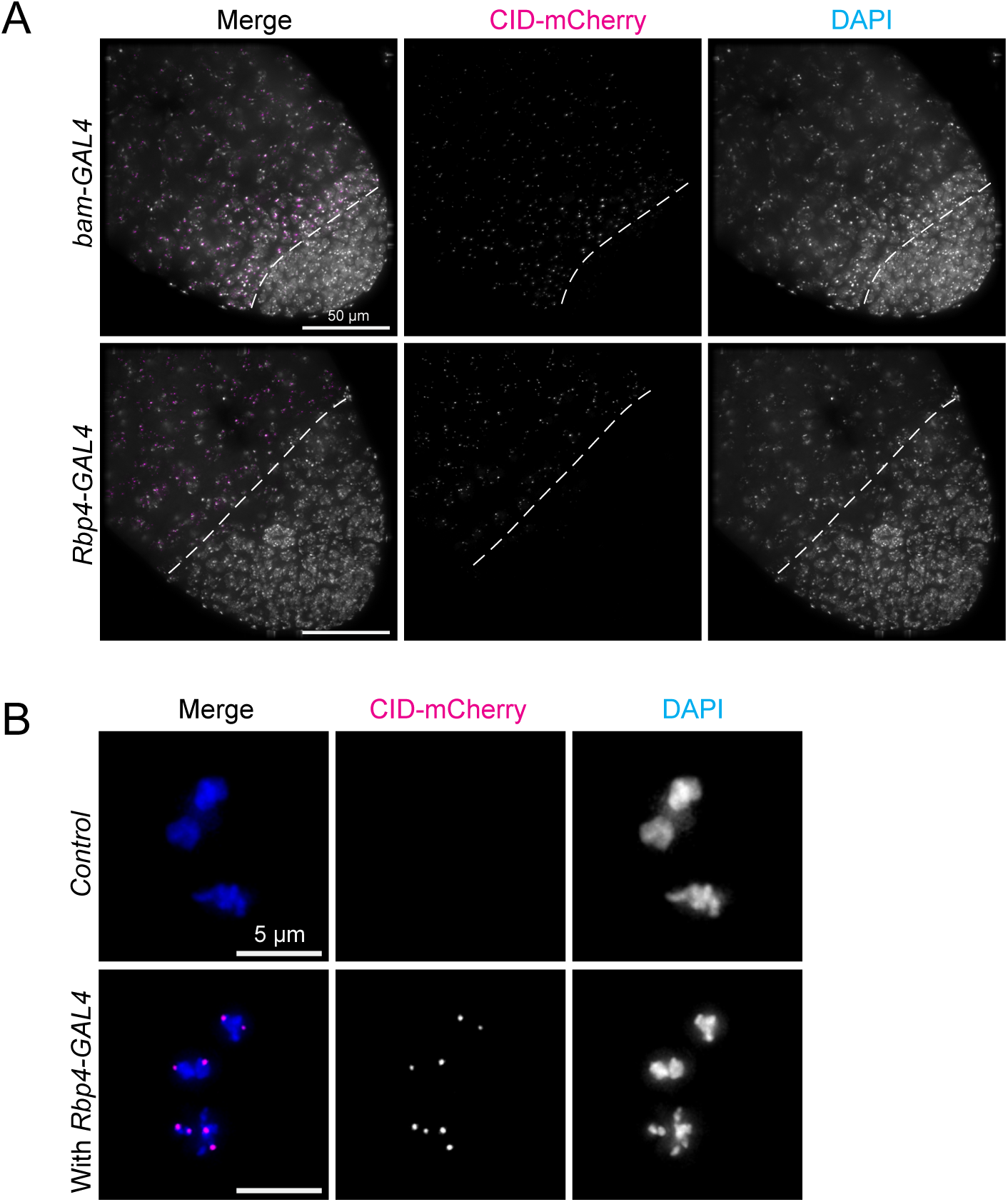
Comparison of *bam-GAL4* and *Rbp4-GAL4* expression timing in *Drosophila* testis. (**A**) Representative images of adult testis apices from crosses of either *bam*-GAL4 or *Rbp4-GAL4* to the UAS-CID-mCherry fly line showing GAL4-driven expression of mCherry tagged CID (magenta) and DAPI (blue). Scale bar = 50 µm. (**B**) Prometaphase I nuclei from pupal testes stained with DAPI (blue), showing *Rbp4-GAL4* driven expression of mCherry tagged CID protein (magenta). The control comprised of the UAS-CID-mCherry line without crossing to *Rbp4-GAL4*. Scale bar = 5 µm.

